# Structural variability of EspG chaperones from mycobacterial ESX-1, ESX-3 and ESX-5 type VII secretion systems

**DOI:** 10.1101/250803

**Authors:** Anne T. Tuukkanen, Diana Freire, Sum Chan, Mark A. Arbing, Robert W. Reed, Timothy J. Evans, Grasilda Zenkeviciutė, Jennifer Kim, Sara Kahng, Michael R. Sawaya, Catherine T. Chaton, Matthias Wilmanns, David Eisenberg, Annabel H. A. Parret, Konstantin V. Korotkov

## Abstract

Type VII secretion systems (ESX) are responsible for transport of multiple proteins in mycobacteria. How different ESX systems achieve specific secretion of cognate substrates remains elusive. In the ESX systems, the cytoplasmic chaperone EspG forms complexes with heterodimeric PE-PPE substrates that are secreted from the cells or remain associated with the cell surface. Here we report the crystal structure of the EspG_1_ chaperone from the ESX-1 system determined using a fusion strategy with T4 lysozyme. EspG_1_ adopts a quasi 2-fold symmetric structure that consists of a central β-sheet and two α-helical bundles. Additionally, we describe the structures of EspG_3_ chaperones from four different crystal forms. Alternate conformations of the putative PE-PPE binding site are revealed by comparison of the available EspG_3_ structures. Analysis of EspG_1_, EspG_3_ and EspG_5_ chaperones using small-angle X-ray scattering (SAXS) reveals that EspG_1_ and EspG_3_ chaperones form dimers in solution, which we observed in several of our crystal forms. Finally, we propose a model of the ESX-3 specific EspG_3-_PE5-PPE4 complex based on the SAXS analysis.

**Highlights:**

- The crystal structure of EspG_1_ reveals the common architecture of the type VII secretion system chaperones
- Structures of EspG_3_ chaperones display a number of conformations that could reflect alternative substrate binding modes
- EspG_3_ chaperones dimerize in solution
- A model of EspG_3_ in complex with its substrate PE-PPE dimer is proposed based on SAXS data

## Introduction

The most deadly bacterial pathogen worldwide is *Mycobacterium tuberculosis* (*Mtb*), which causes tuberculosis (TB). While many infectious diseases can be controlled by vaccination, TB lacks an effective vaccine and even prior infection with *Mtb* does not provide lasting immunity. Moreover, standard anti-TB therapy requires the use of a combination of drugs for six months, which leads to poor compliance and to emergence of drug resistance [1]. Even more threatening is the global increase in extensively drug-resistant *Mtb* and the emergence of extremely drug-resistant *Mtb*. Anti-virulence drugs targeting mycobacterial secretion have the potential to become a valuable alternative to classical antibiotics [2]. During infection, pathogenic mycobacteria use several related protein secretion pathways designated ESX systems [3–5]. The *Mtb* genome encodes five such secretion systems, ESX-1 through ESX-5. Each consists of ATPases, membrane proteins, a protease, accessory proteins, and secreted substrates [6, 7]. Four conserved components of the ESX-5 system — EccB_5_, EccC_5_, EccD_5_ and EccE_5_ — form a platform complex with six-fold symmetry that is embedded in the mycobacterial inner membrane [8, 9]. The ESX-1 core complex is composed of paralogous components [10], which suggests that all ESX systems assemble into similar complexes. Many ESX-secreted substrates are interdependent on each other for secretion, suggesting that they might be a part of the ESX secretion machinery [11, 12]. The most abundant class of ESX substrates is represented by the so-called PE and PPE proteins [13]. These proteins generally form alpha-helical heterodimers that are probably secreted in a folded conformation [14]. Several PE/PPE proteins are major antigens for TB diagnostic and vaccine development [15–19]. Importantly, PE/PPE proteins are secreted specifically by their cognate ESX secretion systems [20–24] raising the question of how various ESX systems discriminate among PE/PPE substrates.

Previously it was demonstrated that PE/PPE protein secretion in *Mycobacterium marinum* is impaired upon disruption of the *espG* gene encoded within its respective ESX gene locus leading to accumulation of substrates in the bacterial cytosol [25]. The crystal structure of the heterotrimeric EspG_5_-PE25–PPE41 protein complex revealed that EspG_5_ interacts with a PE25–PPE41 heterodimer by binding to a hydrophobic patch at the tip of PPE41 [26, 27]. The general YxxD/E secretion motif [28, 29] at the distal end of PE25 is free to interact with the ESX-5 secretion machinery in the inner membrane, probably by interaction with the Ftsk/SpoIIIE-like ATPase EccC_5_ [30, 31]. In addition, EspG_5_ was reported to improve solubility of aggregation-prone PE–PPE pairs upon co-expression [26, 32]. Thus, EspG acts as a disaggregase of ESX substrates in the cytosol prior to secretion. Moreover, substrate specificity is determined by the EspG-binding domain of PPE proteins as demonstrated by substrate re-routing experiments [33]. While structures of the EspG_5_ chaperone in complexes with PE–PPE substrates and a monomeric EspG_3_ chaperone have been reported [26, 27, 34], structural information on EspG_1_ is lacking.

In this study, we report the first crystal structure of an EspG_1_ chaperone from *Mycobacterium kansasii* and four crystal structures of EspG_3_ from *M*. *smegmatis and M. marinum*. We analyze here the available atomic structures of EspG chaperones and present a thorough study of the conformational variability of EspG proteins in apo and substrate-bound forms (EspG-PE–PPE) using small-angle X-ray scattering (SAXS). In addition, we characterize the SAXS-based rigid-body structure of the EspG_3_-PE5–PPE4 protein complex in solution and compare it to the atomic structure of EspG_5_-PE25–PPE41 in order to obtain further insights into protein flexibility and substrate recognition. Our study shows that the EspG chaperones are capable of adopting multiple conformational states, likely a key determinant of their ability to recognize multiple PE–PPE substrates.

## Results

### The crystal structure of *M. kansasii* EspG_1_

Initial attempts to crystallize *M. tuberculosis* EspG_1_ (EspG_1mtu_) were not successful; therefore we screened several homologs of EspG_1_ from other mycobacterial species. We obtained microcrystals using an optimized construct of *M. kansassi* EspG_1_ (EspG_1mka_) that has 80% sequence identity with EspG_1mtu_ (Supplementary Fig. S1). However, extensive optimization of these crystals did not lead to diffraction quality crystals. To overcome these difficulties, we utilized a fusion approach using maltose binding protein or T4 lysozyme (T4L) as the N-terminal fusions. Whereas maltose binding protein fusion did not crystallize, crystals of the T4L-EspG_1mka_ fusion could be readily optimized and diffracted to 2.27 Å resolution. The structure of T4L-EspG_1mka_ was solved by molecular replacement and refined to R_work_ 0.214 and R_free_ 0.251 with good geometry (Table 1). The structure contains two molecules in the asymmetric unit (Fig. 1a) with an extensive interface between the T4L moieties (2250 Å^2^ buried surface area). Surprisingly, part of the TEV cleavage sequence at the N-terminus of T4L is ordered in the structure and contributes both to the T4L dimer interface (849 Å^2^ buried surface area corresponding to 38% of the T4L–T4L interface) and the intra-subunit contacts between T4L and EspG_1mka_. The conformations of the two copies of EspG_1mka_ in the asymmetric unit are very similar and superimpose with a root-mean-square deviation (RMSD) of 1.0 Å over 231 Cα atoms (Supplementary Fig. S2). EspG_1mka_ has a typical EspG fold characterized by a central anti-parallel β-sheet and two α-helical bundles (Fig. 1b). Several parts of the EspG_1mka_ structure did not have interpretable electron density and were not modeled, including the loop preceding the α2 helix (chains A and B), the β2–β3 loop (chain A), part of the β6 strand (chain B), and the α6 helix (chains A and B) (Fig. 1a). The residues corresponding to the C-terminal helical bundle displayed higher *B* factors compared to other parts of the structure (Supplementary Fig. S2b). The C-terminal helical bundle likely forms part of the substrate recognition site and could become more ordered upon binding of a cognate PE–PPE dimer. The N-terminal and C-terminal subdomains of EspG_1mka_ are related by a quasi two-fold symmetry, have 10% sequence identity, and can be superimposed with a RMSD of 2.7 Å over 71 Cα atoms (Supplementary Fig. S3).

**Table 1.** Data collection and refinement statistics.

|  | T4L-EspG<sub>1</sub> | EspG<sub>3</sub> | EspG<sub>3</sub> | EspG<sub>3</sub> | EspG<sub>3</sub> |
| --- | --- | --- | --- | --- | --- |
|  | *M. kansasii* | *M. marinum* | *M. smegmatis* | *M. smegmatis* | *M. smegmatis* |
|  | PDB 5VBA | PDB 5DLB | PDB 4L4W | PDB 4RCL | PDB 5SXL |
| **Data collection** |  |  |  |  |  |
| Wavelength (Å) | 1.0000 | 1.0702 | 0.9793 | 1.0000 | 1.0000 |
| Space group | *P*2<sub>1</sub>2<sub>1</sub>2<sub>1</sub> | *C*2 | *C*222<sub>1</sub> | *P*4<sub>3</sub>2<sub>1</sub>2 | *P*3<sub>2</sub>21 |
| Cell dimensions: |  |  |  |  |  |
| *a*, *b*, *c* (Å) | 64.14, 81.69,<br>160.02 | 112.21, 46.02,<br>58.01 | 47.08, 123.69,<br>180.85 | 92.68, 92.68, 158.85 | 59.15, 59.15,<br>183.1 |
| $\alpha$, $\beta$, $\gamma$ (°) | 90, 90, 90 | 90, 92.24, 90 | 90, 90, 90 | 90, 90, 90 | 90, 90, 120 |
| Resolution (Å) | 59.54–2.27 (2.33–<br>2.27)<sup>a</sup> | 57.96–1.77 (1.87–<br>1.77) | 45.21–2.04 (2.09–<br>2.04) | 46.34–2.70 (2.77–2.70) | 39.24–2.46<br>(2.52–2.46) |
| *R*<sub>sym</sub> | 0.097 (1.110) | 0.066 (0.827) | 0.087 (0.809) | 0.087 (1.600) | 0.075 (0.882) |
| CC<sub>1/2</sub><sup>b</sup> | 99.7 (58.3) | 99.5 (65.9) | 99.7 (64.6) | 99.9 (76.2) | 99.8 (79.3) |
| *I* / σ*I* | 11.1 (1.74) | 12.6 (1.40) | 12.2 (1.55) | 19.8 (1.92) | 12.2 (1.62) |
| Completeness (%) | 99.8 (100.0) | 89.0 (90.0) | 97.6 (79.5) | 99.8 (98.5) | 98.6 (99.8) |
| Multiplicity | 6.3 (6.5) | 3.2 (3.4) | 5.5 (3.1) | 11.1 (11.2) | 5.4 (5.5) |
| **Refinement** |  |  |  |  |  |
| Resolution (Å) | 59.54–2.27 | 57.96–1.77 | 45.21–2.04 | 46.34–2.70 | 39.24–2.46 |
| No. reflections (total / free) | 39576 / 1882 | 24426 / 1310 | 33548 / 1682 | 19765 / 1008 | 13963 / 707 |
| $R_{\text{work}} / R_{\text{free}}$ | 0.214 / 0.251 | 0.209 / 0.245 | 0.186 / 0.231 | 0.230 / 0.278 | 0.206 / 0.247 |
| Number of atoms: |  |  |  |  |  |
| Protein | 6288 | 1944 | 4003 | 3709 | 2078 |
| Ligand/ion | 10 | 140 |  |  |  |
| Water | 56 | 65 | 225 | 14 | 17 |
| *B*-factors: |  |  |  |  |  |
| Protein | 61.6 | 37.8 | 37.9 | 92.1 | 81.7 |
| Ligand/ion | 68.1 | 72.7 |  |  |  |
| Water | 48.5 | 47.1 | 39.3 | 40.6 | 63.4 |
| All atoms | 61.5 | 40.4 | 38.0 | 91.9 | 81.6 |
| Wilson *B* | 47.8 | 38.0 | 37.2 | 74.7 | 67.6 |
| R.m.s. deviations: |  |  |  |  |  |
| Bond lengths (Å) | 0.004 | 0.019 | 0.008 | 0.003 | 0.002 |
| Bond angles (°) | 0.657 | 2.130 | 1.166 | 0.722 | 0.494 |
| Ramachandran distribution<sup>c</sup> |  |  |  |  |  |
| (%): |  |  |  |  |  |
| Favored | 97.73 | 94.16 | 97.32 | 96.07 | 95.97 |
| Outliers | 0.13 | 0.39 | 0 | 0.21 | 0.37 |

<sup>a</sup> Values in parentheses are for the highest-resolution shell.
<sup>b</sup> CC<sub>1/2</sub> correlation coefficient as defined in Karplus & Diederichs [42] and calculated by *XSCALE* [41].
<sup>c</sup> Calculated using the MolProbity server (http://molprobity.biochem.duke.edu) [76].

**Figure 1.**
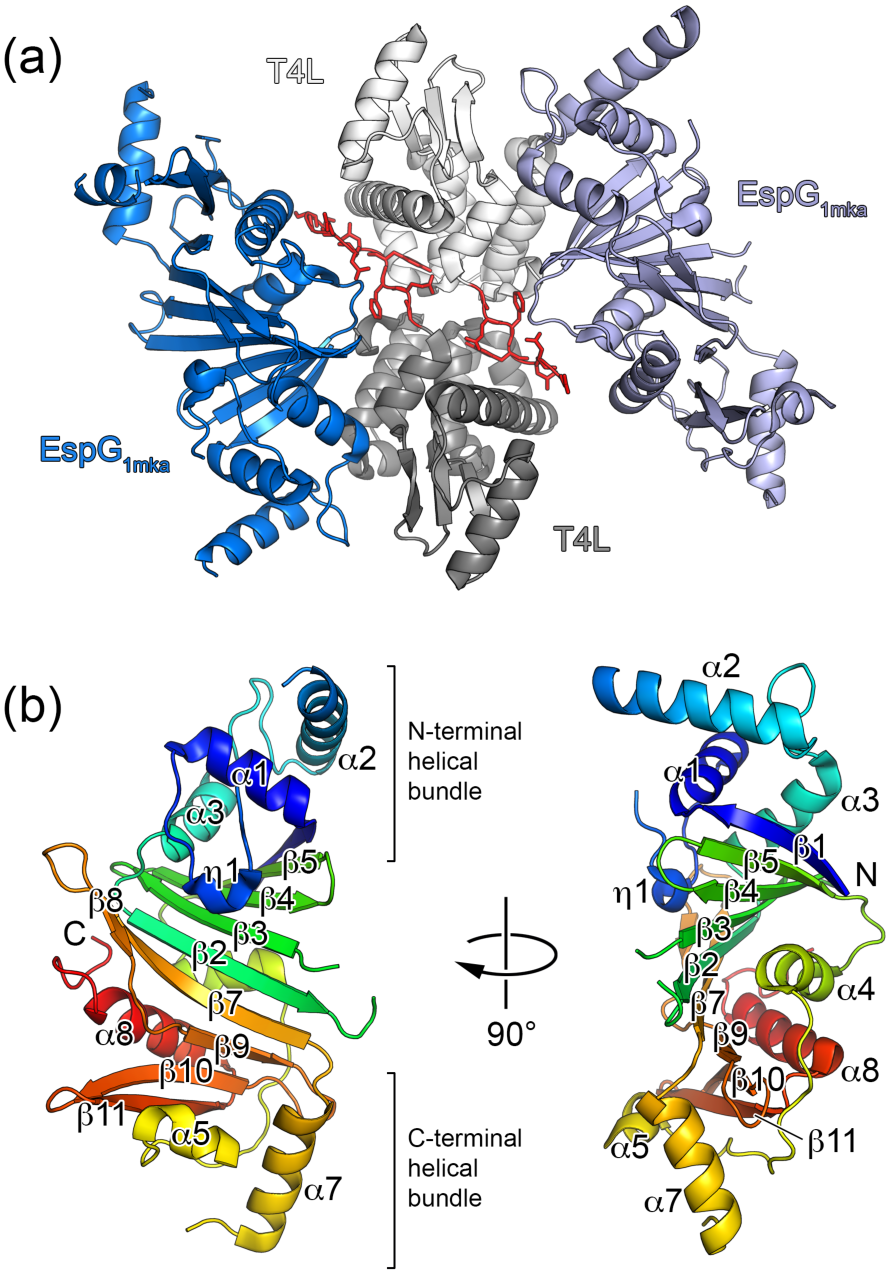
Crystal structure of *M. kansasii* EspG_1_ (EspG_1mka_). (a) View of the two subunits of the T4L-EspG_1mka_ fusion protein in the asymmetric unit. Chain A is shown in light grey (T4L) and light blue (EspG_1mka_), and chain B is shown in dark grey (T4L) and dark blue (EspG_1mka_). Residues corresponding to the TEV cleavage sequence are shown in red with side chains in stick representation. (b) A monomer of EspG_1mka_ is shown in ribbon representation colored in rainbow colors from N-terminus (blue) to C-terminus (red). The N-terminal T4L fusion moiety is not shown for clarity.

### The EspG fold is conserved in ESX-1, ESX-3 and ESX-5 systems

In order to extend the structural knowledge of the EspG-substrate interaction and the differences between these interactions in ESX-1, ESX-3 and ESX-5 secretion, we determined additional crystal structures of the EspG chaperones from the ESX-1 and ESX-3 systems (Table 1). Despite the fact that EspG chaperones display the lowest level of protein sequence similarity (13–23% sequence identity) of all the core components of the ESX systems, the EspG_1_, EspG_3_ and EspG_5_ structures have a highly similar fold (Fig. 2, Fig. 3, Supplementary Figs. S4 and S5 and Supplementary Table S1). RMSD of the aligned atoms for EspG superposition is 2.4 Å for EspG_1mka_ vs. EspG_5mtu_ over 224 Cα atoms, 2.4 Å for EspG_3mma_ vs. EspG_5mtu_ over 238 Cα atoms and 2.4 Å for EspG_1mka_ vs. EspG_3mma_ over 234 Cα atoms. Despite the high overall structural similarity, the C-terminal helical bundles of EspG_1mka_ and EspG_3_ structures have a distinct conformation, when compared to the EspG_5mtu_ structure bound to the PE25–PPE41 dimer, which appears to be incompatible with substrate binding (Fig. 3). Another significant difference is the length and the conformation of the β2–β3 loop. In the EspG_5mtu_-PE25–PPE41–structure (PDB ID 4KXR [26]), it extends 23 amino acid residues (Gly^92^-Asn^114^) and interacts strongly with the PE25–PPE41 dimer, whereas, for example, in the EspG_3msm_ (PDB ID 4L4W) structure the loop consists of only 12 amino acid residues (Ser^87^-Leu^98^). These structural differences could be explained by conformational changes in EspG_5mtu_ induced by PE–PPE binding, suggesting that binding of EspG_1_ and EspG_3_ to their cognate PPE protein partners is different from the EspG_5_-PPE41 interaction observed in the EspG_5mtu_-PE25–PPE41–structure.

**Figure 2.**
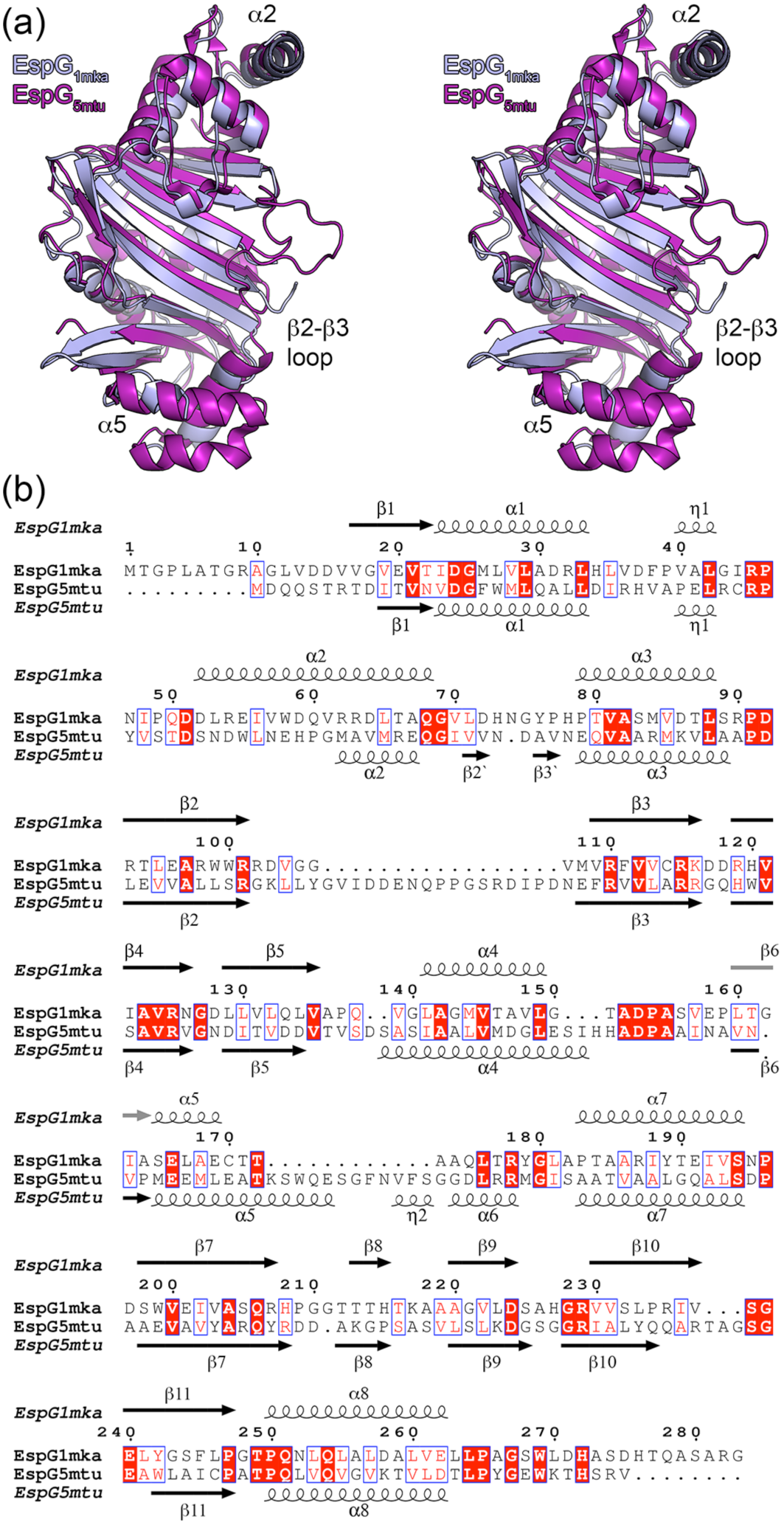
Structural comparison between EspG_1mka_ and EspG_5mtu_. (a) Stereo view of superposed EspG_1mka_ and EspG_5mtu_ crystal structures. The structure of the EspG_5mtu_ monomer is derived from the trimeric EspG_5mtu_-PE25–PPE41–complex (PDB ID 4KXR, [26]). (b) Structure-based sequence alignment of EspG_1mka_ and EspG_5mtu_. Secondary structure elements corresponding to the EspG_1mka_ structure (PDB ID 5VBA) and EspG_5mtu_ structure (PDB ID 4KXR) are displayed above and below the alignment.

**Figure 3.**
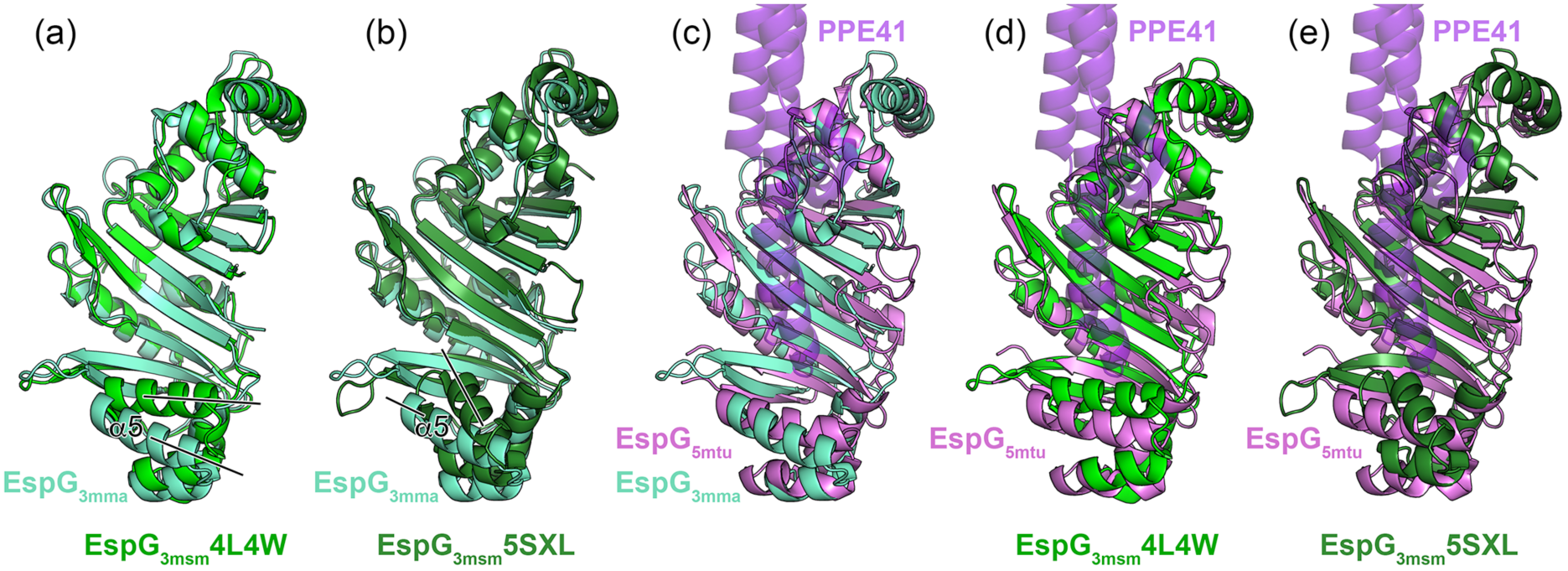
Crystal structures of EspG_3_ chaperones display variations in their putative PE-PPE binding region. (a) Structural superposition of EspG_3mma_ (aquamarine) and EspG_3msm_ (PDB ID 4L4W, green). Black lines indicate differences in the orientation of the α5 helix. A stereo version is available as Supplementary Figure 4a. (b) Structural superposition of EspG_3mma_ and EspG_3msm_ (PDB ID 5SXL, dark green). A stereo version is available as Supplementary Figure 4b. (c,d,e) Structural superposition of EspG_3mma_, EspG_3msm_ (PDB ID 4L4W), and EspG_3msm_ (PDB ID 5SXL) with EspG_5mtu_ (PDB ID 4W4I [27], violet) derived from the heterotrimeric EspG_5mtu_-PE25-PPE41 structure (PDB ID 4KXR [26]). PPE41 (purple) is shown in semi-transparent ribbon representation. PE25 is omitted for clarity as it is not in contact with EspG_5mtu_. Stereo versions of (c,d,e) are available as Supplementary Figure 5.

### Variation of quaternary structure within the EspG protein family

Analysis of crystal packing in the available EspG_3_ structures revealed a number of possible quaternary arrangements in addition to the monomeric state. Firstly, a “wing-shaped dimer” was found in the asymmetric unit of the EspG_3msm_ (PDB ID 4L4W) structure with 1892 Å^2^ buried surface area (Fig. 4). The dimeric interface is mediated by residues from the C-terminal helical bundles and strands β6, β10, β11 and helix α8. In contrast, the asymmetric unit of the EspG_3msm_ (PDB ID 4RCL) structure contains a dimer in front-to-front orientation with 1761 Å^2^ buried surface area. The β8 strands from the two subunits are located at the core of the interface and form an inter-subunit β-sheet. This dimeric conformation is further referred to as a “β8-mediated dimer”. However, in addition a “wing-shaped dimer” similar to the EspG_3msm_ (PDB ID 4L4W) structure is present in the crystal lattice, with 2054 Å^2^ buried surface area. Furthermore, the asymmetric unit of the EspG_3msm_ (PDB ID 4W4J [27]) structure also contains a similar wing-shaped dimer with substantial buried surface area as well as an β8-mediated dimer (Fig. 4).

**Figure 4.**
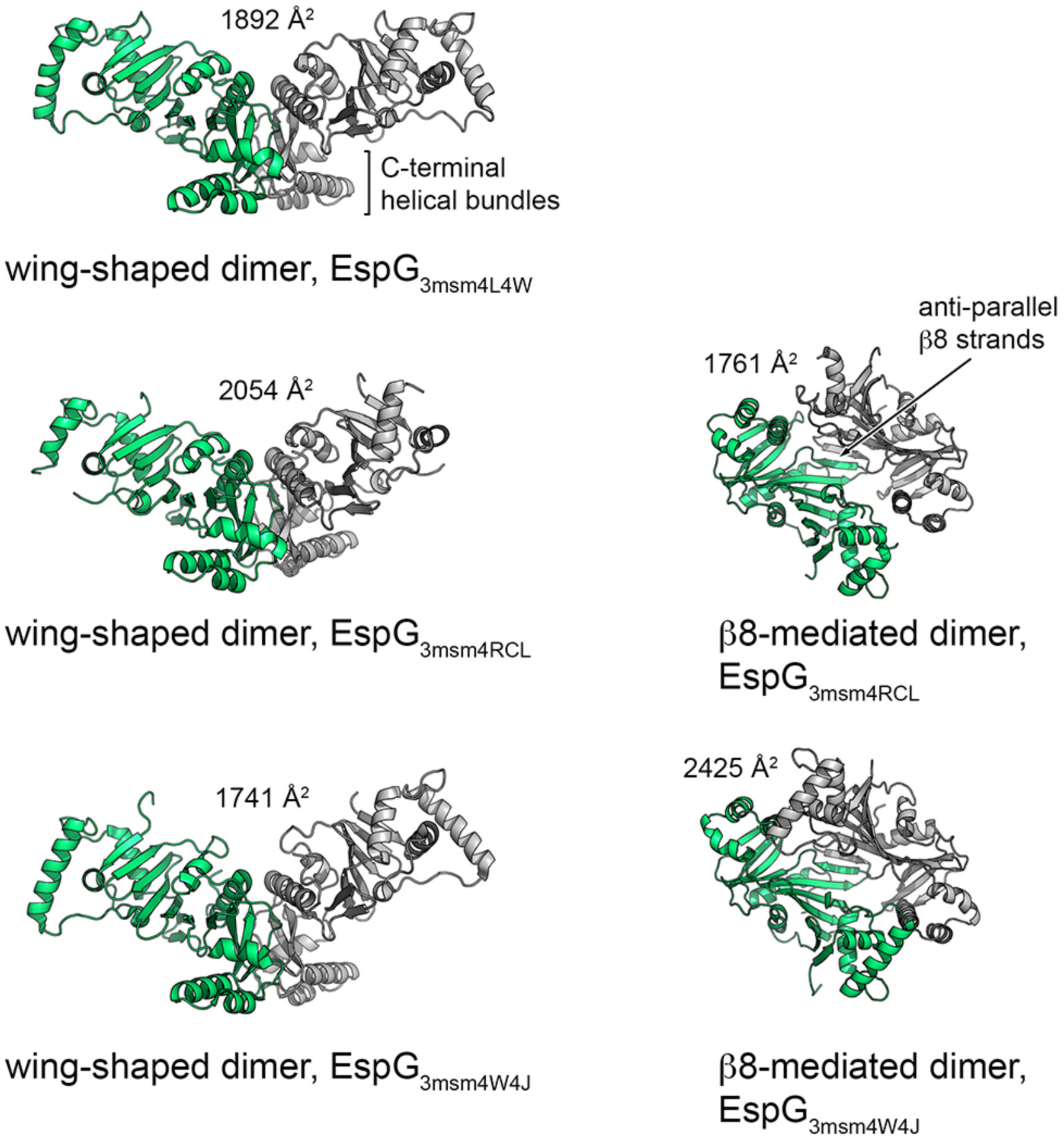
Cartoon representation of the common dimer structures observed in crystal forms of EspG_3_. Superimposed subunits are in green with the buried surface area of the dimer interface indicated above the structure. The wing-shaped dimers are present in the asymmetric unit of EspG_3msm_ (PDB ID 4L4W) and EspG_3msm_ (PDB ID 4W4J) or generated by crystallographic symmetry in EspG_3msm_ (PDB ID 4RCL). The β8-mediated dimer is present in the asymmetric unit in EspG_3msm_ (PDB ID 4RCL) and generated by crystallographic symmetry in EspG_3msm_ (PDB ID 4W4J [27]).

Altogether, EspG_3msm_ wing-shaped dimers are observed in three independent crystal structures, and β8-mediated dimers are seen in two crystal structures. The dimer interfaces are highly similar, with the subunits rotated relative to each other by 12 degrees in the wing-shaped dimers and 5 degrees in the β8-mediated dimers (Supplementary Fig. S6). The different quaternary structures of EspG chaperones reflect the variability that exists within this protein family (Table 2). As previously proposed [26], EspG likely acts as a chaperone that maintains PE/PPE secretion targets in the cytosol in a soluble state. Further experiments will be required to elucidate whether the dimerization of EspG_3msm_ plays a role in the function of the chaperone *in vivo*. To assess whether EspG proteins of ESX-1 and ESX-5 display similar oligomerization behavior in solution, we performed a structural analysis of several EspG proteins in solution.

**Table 2.**
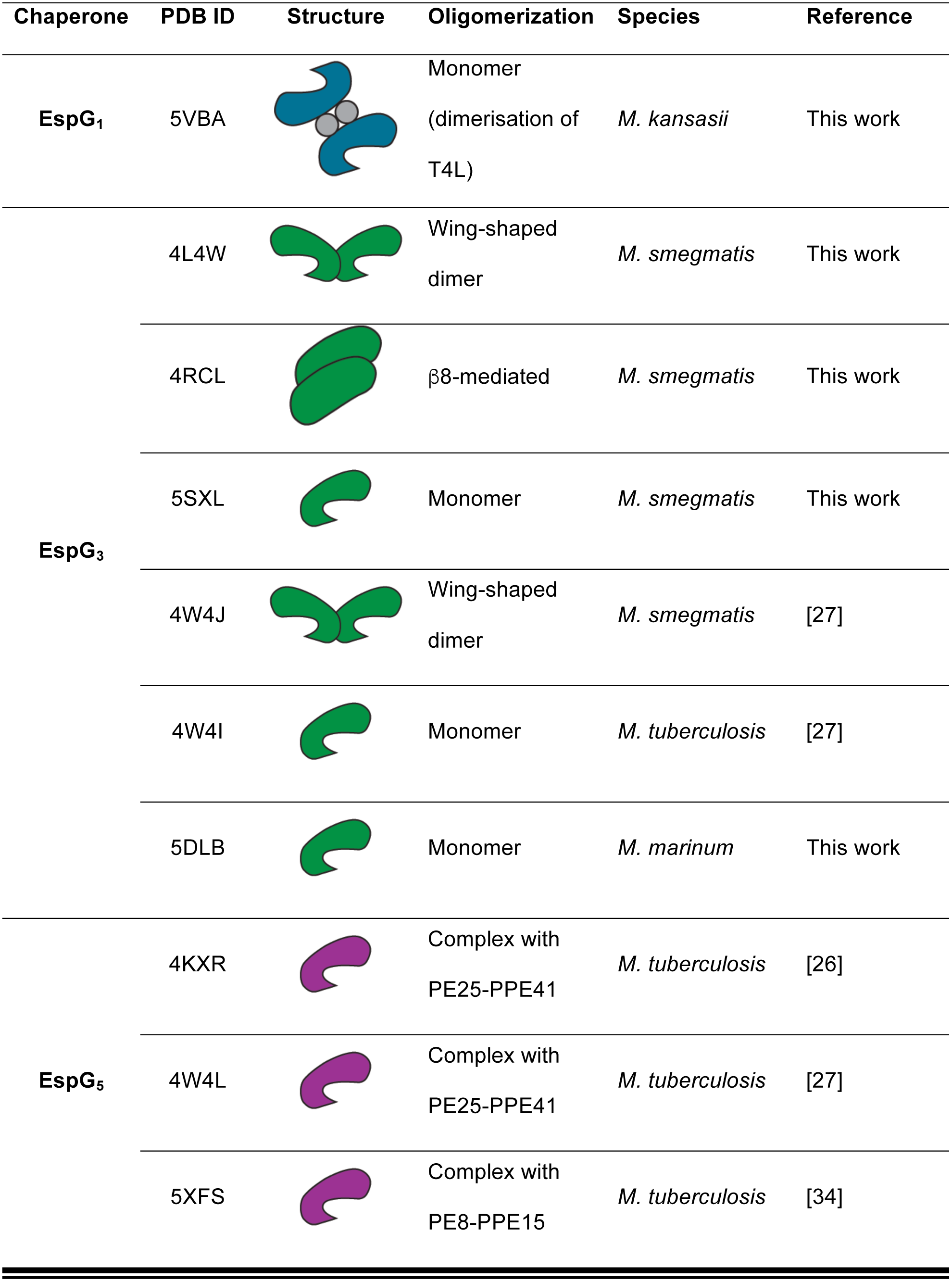
Overview of EspG crystal structures.

### Concentration-dependent oligomerization of EspG chaperones

We studied solution structures of EspG proteins from ESX-1 (EspG_1mma_), ESX-3 (EspG_3mma_, EspG_3mtu_, EspG_3msm_) and ESX-5 (EspG_5mtu_) secretion systems using SAXS (Table 3). The EspG proteins from different secretion systems have diverged significantly although the EspG_3_ homologs from different species have a high degree of sequence identity. The Guinier analysis of the obtained SAXS profiles confirmed that the proteins were not aggregated allowing further analysis of their structures and oligomeric states (Table 3). The dependencies of the molecular weight estimates and the excluded volume of the hydrated particles on concentration were studied for all proteins. Prior structural information of the monomeric form existed for all of the studied proteins, but experimental models of the dimeric state were only available for EspG_3msm_.

**Table 3.** Collected SAXS data

| Data collection parameters |  |  |  |  |  |  |  |  |  |
| --- | --- | --- | --- | --- | --- | --- | --- | --- | --- |
| Instrument |  | P12 at EMBL/DESY, storage ring PETRA III, Germany |  |  |  |  |  |  |  |
| Beam geometry (mm<sup>2</sup>) |  | 0.2 × 0.12 |  |  |  |  |  |  |  |
| Wavelength (Å) |  | 1.24 |  |  |  |  |  |  |  |
| q-range (Å<sup>-1</sup>) |  | 0.0023 – 0.47 |  |  |  |  |  |  |  |
| Exposure time (ms) |  | 20 × 50 |  |  |  |  |  |  |  |
| Temperature (K) |  | 283 |  |  |  |  |  |  |  |
| Instrument |  | B21 at Diamond Light Source, United kingdom |  |  |  |  |  |  |  |
| Beam geometry (mm<sup>2</sup>) |  | 1.0 x 5.0 |  |  |  |  |  |  |  |
| Wavelength (Å) |  | 1.0 |  |  |  |  |  |  |  |
| q-range (Å<sup>-1</sup>) |  | 0.0038 – 0.42 |  |  |  |  |  |  |  |
| Exposure time (ms) |  | 60 x 100 |  |  |  |  |  |  |  |
| Temperature (K) |  | 283 |  |  |  |  |  |  |  |
| Structural parameters |  |  |  |  |  |  |  |  |  |
| Sample | Conc.<br>(mg ml<sup>-1</sup>) | *R*<sub>g, Guinier</sub><br>(nm) | *R*<sub>g, Pr</sub><br>(nm) | *D*<sub>max</sub><br>(nm) | *MM*<sub>theor</sub><br>(kDa) | *MM*<sub>SAXS</sub><br>(kDa) | *MM*<sub>POROD</sub><br>(kDa) | *MM*<sub>DAM</sub><br>(kDa) | *Ab initio*<br>resolution,<br>(Å) |
| EspG<sub>1mma</sub> | 1.0 | 2.7 | 2.9 | 9.7 | 29.8 | 25.0 | 43.0 | 45.0 | 42 ± 3 |
|  | 2.0 | 3.2 | 3.1 | 11.0 | 29.8 | 32.0 | 55.0 |  |  |
|  | 4.2 | 3.4 | 3.5 | 11.0 | 29.8 | 37.0 | 65.0 |  |  |
|  | 6.0 | 3.5 | 3.6 | 12.0 | 29.8 | 40.0 | 72.0 |  |  |
| EspG<sub>3mtu</sub> | 1.1 | 2.5 | 2.6 | 9.0 | 31.6 | 23.0 | 41.0 | 41.0 | 34 ± 3 |
|  | 2.0 | 2.8 | 3.0 | 10.0 | 31.6 | 30.0 | 47.0 |  |  |
|  | 4.0 | 3.2 | 3.1 | 10.0 | 31.6 | 32.0 | 56.0 |  |  |
|  | 6.1 | 3.5 | 3.5 | 12.0 | 31.6 | 39.0 | 68.0 |  |  |
| EspG<sub>3mma</sub> | 1.1 | 2.3 | 2.5 | 8.0 | 32.0 | 20.0 | 39.0 | 34.0 | 23 ± 2 |
|  | 2.1 | 2.3 | 2.4 | 8.0 | 32.0 | 22.0 | 35.0 | 35.0 |  |
|  | 4.0 | 2.3 | 2.4 | 8.0 | 32.0 | 21.0 | 40.0 | 38.0 |  |
|  | 6.0 | 2.3 | 2.3 | 8.0 | 32.0 | 22.0 | 44.0 | 36.0 |  |
| EspG<sub>3msm</sub> | 0.8 | 2.5 | 2.6 | 8.6 | 31.6 | 20.5 | 39.0 | 48.0 | 36 ± 3 |
|  | 1.8 | 2.5 | 2.5 | 8.8 | 31.6 | 21.4 | 32.0 | 41.0 |  |
|  | 3.9 | 2.6 | 2.6 | 9.3 | 31.6 | 22.1 | 39.0 | 44.0 |  |
|  | 6.2 | 2.8 | 2.7 | 9.7 | 31.6 | 24.0 | 39.0 | 47.0 |  |
| EspG<sub>3msm</sub><br>Se-Met | 0.9 | 2.5 | 2.5 | 9.2 | 31.6 | 22.5 | 38.0 | 43.0 | 39 ± 3 |
|  | 2.1 | 2.5 | 2.6 | 9.2 | 31.6 | 22.5 | 38.0 |  |  |
|  | 3.8 | 2.6 | 2.6 | 9.2 | 31.6 | 24.3 | 40.0 |  |  |
| EspG<sub>5mtu</sub> | 0.9 | 2.3 | 2.5 | 8.0 | 32.4 | 23.0 | 41.0 | 47.0 | 25 ± 2 |
|  | 1.7 | 2.4 | 2.4 | 8.0 | 32.4 | 22.0 | 42.0 |  |  |
|  | 4.1 | 2.4 | 2.5 | 8.0 | 32.4 | 21.0 | 43.0 |  |  |
|  | 6.7 | 2.4 | 2.4 | 8.0 | 32.4 | 22.0 | 41.0 |  |  |
| EspG<sub>5mtu</sub>-<br>PE25-PPE41 | 0.5 | 4.0 | 4.0 | 13.0 | 65.0 | 76.0 | 46.0 | 76.0 | 37 ± 3 |
| EspG<sub>3msm</sub>-<br>PE5-PPE4 | 1.2 | 4.0 | 4.2 | 14.2 | 95.0 | 83.0 | 52.0 | 83.0 | 38 ± 3 |

SAXS measurements of EspG_1mma_ at four different protein concentrations resulted in distinct scattering profiles. The molecular weight (based on the Porod volume) increased with increasing concentration from 43 to 72 kDa at 1.0 mg/mL and 6.0 mg/mL, respectively (Table 3). A similar trend was observed for the *R_g_* and *D_max_* values. The oligomer analysis of the concentration-dependent SAXS data was carried out using theoretical scattering profiles based on the monomeric EspG_1mka_ (PDB ID 5VBA) structure and two different dimeric structural models. The β8-mediated dimer observed in the EspG_3msm_ (PDB ID 4RCL) structure and the wing-shaped dimer structure of EspG_3msm_ (PDB ID 4L4W) were used as templates to generate EspG_1mma_ dimer models. The corresponding theoretical scattering profiles were exploited to decompose the experimental data (Table 4). The *OLIGOMER* analysis showed that EspG_1mma_ is predominantly a monomer at low protein concentrations, while the fraction of dimeric protein increases up to 50% at the highest concentration measured. However, the goodness-of-fit (χ^2^) of the theoretical scattering based on linear combinations of theoretical monomer/dimer scattering profiles to the experimental SAXS data varied significantly between the two dimer models (Table 4). The SAXS data could not be interpreted successfully using the β8-mediated dimeric arrangement (Table 4). Additional evidence that the β8-mediated dimer is a non-physiological crystallographic dimer is the substantial structural clashes observed when the β8-mediated EspG_1_ dimer is superimposed onto the EspG_5mtu_ structure derived from the heterotrimeric EspG_5mtu_-PE25–PPE41 crystal structure (Fig. 3). The wing-shaped dimer conformation based on the EspG_3msm_ (PDB ID 4L4W) crystal structure yields better fits (χ^2^ –values between 0.79 and 1.27, Table 4) to the measured scattering data, which strongly indicates that EspG_1_ dimers adopt the wing-shaped conformation in solution.

**Table 4.** Oligomer analysis using the crystallographic monomer and dimer structures of EspG proteins.

| Sample | Template (PDB ID) | Concentration (mg ml<sup>-1</sup>) | Monomeric fraction | Dimeric fraction | Goodness-of-fit, $\chi^2$ |
| --- | --- | --- | --- | --- | --- |
| EspG<sub>1mma</sub> | β8-mediated dimer (4RCL) | 1.0 | 0.94 | 0.06 | 0.80 |
|  |  | 2.0 | 0.74 | 0.26 | 0.93 |
|  |  | 4.2 | 0.58 | 0.42 | 1.44 |
|  |  | 6.0 | 0.50 | 0.50 | 1.90 |
|  | Wing-shaped dimer (4L4W) | 1.0 | 0.93 | 0.07 | 0.79 |
|  |  | 2.0 | 0.72 | 0.28 | 0.86 |
|  |  | 4.2 | 0.54 | 0.46 | 1.01 |
|  |  | 6.0 | 0.43 | 0.57 | 1.27 |
| EspG<sub>3mtu</sub> | Wing-shaped dimer (4L4W) | 1.1 | 0.537 | 0.462 | 0.89 |
|  |  | 2.0 | 0.364 | 0.636 | 0.90 |
|  |  | 4.0 | 0.113 | 0.887 | 0.95 |
|  |  | 6.1 | 0.0 | 0.999 | 0.89 |
| EspG<sub>3mma</sub> | Wing-shaped dimer (4L4W) | 1.1 | 0.531 | 0.468 | 0.82 |
|  |  | 2.1 | 0.195 | 0.805 | 0.88 |
|  |  | 4.0 | 0.000 | 1.000 | 1.05 |
|  |  | 6.0 | 0.00 | 1.000 | 1.69 |
| EspG<sub>3msm</sub> (Native) | Wing-shaped dimer (4L4W) | 0.78 | 0.589 | 0.410 | 0.90 |
|  |  | 1.75 | 0.654 | 0.346 | 1.32 |
|  |  | 3.94 | 0.575 | 0.425 | 1.35 |
|  |  | 6.24 | 0.449 | 0.550 | 2.03 |
| EspG<sub>3msm</sub> (SeMet) | Wing-shaped dimer (4L4W) | 0.90 | 0.70 | 0.30 | 0.92 |
|  |  | 2.14 | 0.70 | 0.30 | 1.80 |
|  |  | 3.79 | 0.59 | 0.41 | 2.50 |

The program *OLIGOMER* was also used to fit a set of theoretical scattering profiles of monomeric and dimeric EspG_3_ models to the experimental EspG_3mtu_ SAXS data. A dimer model was constructed based on the wing-shaped dimer structure of EspG_3msm_ (PDB ID 4L4W). All fits at different concentrations provided good χ^2^ values indicating that our wing-shaped dimer EspG_3mtu_ model based on the EspG_3msm_ structure is appropriate. Analysis of the volume distributions of the oligomeric states showed that even at the lowest concentration (1.1 mg/mL), 46% of the protein was in dimeric form. The dimeric protein fraction increased to 100% at the highest measured concentration (6.1 mg/mL) (Table 4).

Likewise, the volume fractions of the oligomeric states were calculated for EspG_3mma_. Again the dimeric experimental structure of EspG_3msm_ (PDB ID 4L4W) was used as the template to model the dimeric form of EspG_3mma_. EspG_3mma_ showed a similar oligomerization pattern as EspG_3mtu._ The volume fraction of the dimeric form of the protein is above 46% even at the lowest protein concentration (Table 4).

We measured the SAXS profiles of two different EspG_3msm_ protein preparations, native and selenomethionine (SeMet) incorporated forms. Interestingly, the two proteins showed distinct oligomerization behaviors. The native EspG_3msm_ sample had a significant fraction of the dimeric form present already at the lowest measured concentration and the fraction stayed stable as the protein concentration increased (Table 4). On the contrary, the SeMet-labeled form remained monomeric over the whole concentration range (Table 4). The *CRYSOL* fit of the monomeric structure (PDB ID 4L4W) to the scattering data from the SeMet-labeled construct provided a X^2^ value of 0.93. This enabled us to use the SAXS data from the SeMet-labeled protein for *ab initio* modeling, which requires a monodisperse sample (Supplementary Fig. S7a). The SAXS-based *ab initio* model calculated with *DAMMIF* and the monomeric structure of SeMet-labeled EspG_3msm_ (PDB ID 4L4W) are in a good agreement (normalized spatial discrepancy (NSD) = 1.1).

The SAXS profiles of the native form of EspG_5mtu_, the only member of the EspG_5_ family that was analyzed, do not show any concentration dependence over the measured range (0.9 to 6.7 mg/mL) (Table 3). The constant molecular mass estimates suggest that EspG_5mtu_ is monomeric in solution. We used the program *CRYSOL* to fit the theoretical scattering profile based on the crystal structure of the EspG_5mtu_ monomer from the EspG_5mtu_–PE25–PPE41 structure (PDB ID 4KXR) to the experimental SAXS data. A significant misfit was observed in the range of momentum transfer of 1.8 - 2.0 nm^-1^ (Fig. 5a). In order to evaluate whether flexibility of the EspG_5mtu_ β2–β3 loop (Gly^92^-Asn^114^) might cause this discrepancy, we employed molecular dynamics (MD) simulations to assign flexible amino acid residues and ensemble optimization method (EOM) to fit the measured SAXS data. The amino acid residue Root-Mean-Square-Fluctuations (RMSF) monitored during a 2 ns production run indicated high flexibility of the β2–β3 loop region and other shorter loop segments (Fig. 5b). Thus, we introduced flexibility in all loop regions of EspG_5_ with the program EOM and also modeled the 24 amino acid residues that were missing from the crystallographic structure (Fig. 5c). In the EOM approach, the scattering profile is fitted by a linear combination of scattering profiles from several structural models co-existing in solution. The resulting EOM models where the flexible β2–β3 loop exhibits the largest conformational changes compared to the crystallographic structure provide an excellent fit with the experimental SAXS data (Fig. 5a) (χ^2^ = 0.99). In contrast to the original crystallographic structure in which the β2–β3 loop interacts with the PE–PPE heterodimer in an extended conformation, all the EOM structures of EspG_5mtu_ in solution have more compact forms. More specifically, the β2–β3 loop of EspG_5mtu_ folds closely onto the protein core in the SAXS-refined solution structures (Fig. 5c). In addition, the monodisperse EspG_5mtu_ SAXS data were employed for *ab initio* modeling using *DAMMIF* (Supplementary Fig. S7b). The structural alignment yielded a NSD value of 0.93 indicating excellent agreement.

**Figure 5.**
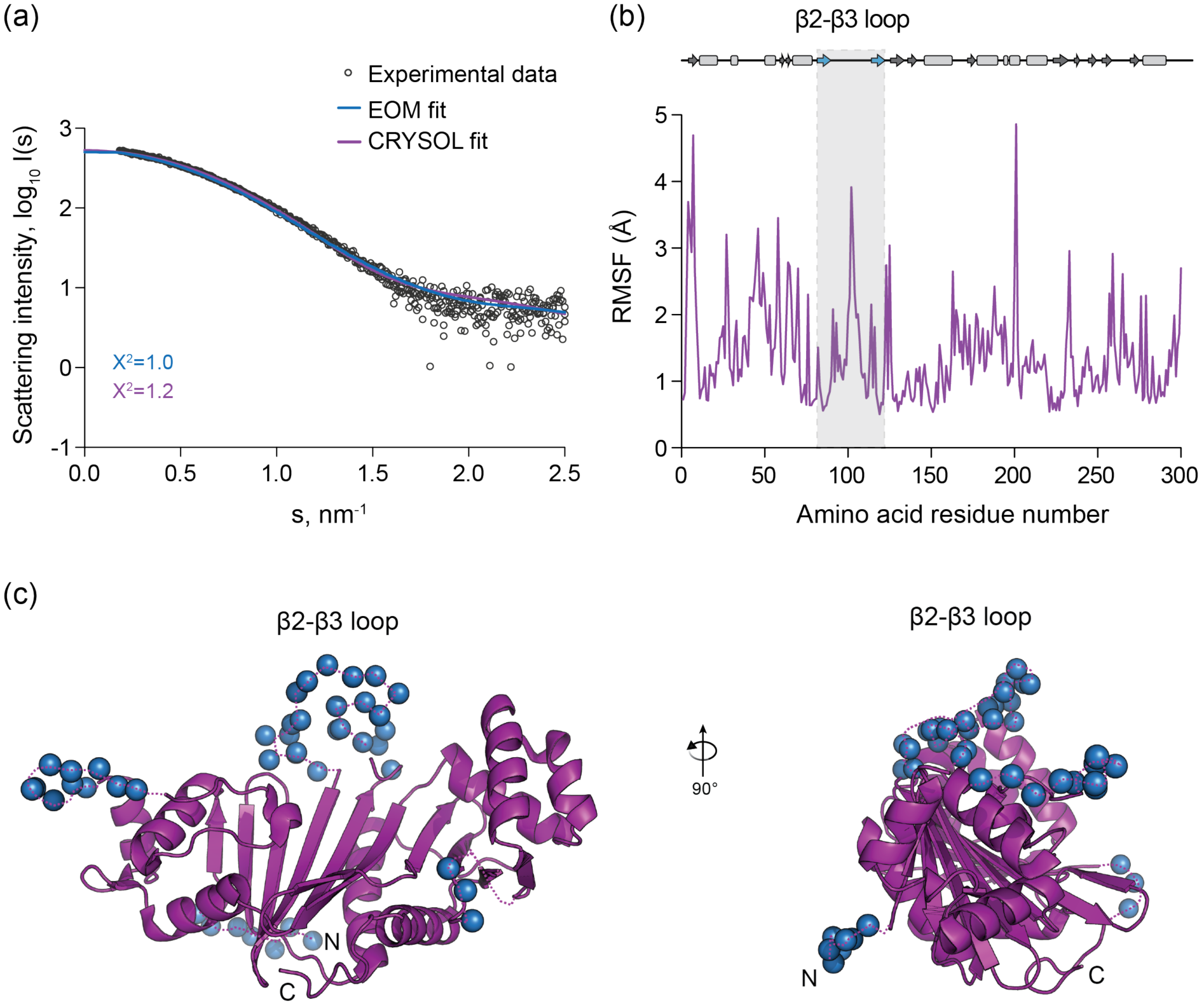
Solution structure of EspG_5mtu_. (a) Experimental SAXS data (black circles) and computed fits (solid lines) with respective discrepancy values (*X^2^)*. The theoretical SAXS curve of the EspG_5mtu_ crystal structure when in complex with PE25/PPE41 as calculated using CRYSOL (purple) (PDB ID 4KXR, chain C) fits the experimental SAXS data with a *X^2^* of 1.2 and an apparent misfit around *s* = 1.8 – 2.0 nm^-1^. To calculate the EOM fit, the 24 amino acid residues that were missing from the crystallographic structure of EspG_5mtu_ were added at the N-terminus and flexibility of all loops was allowed. This procedure resulted in an improvement of the fit to a *X^2^* value of 0.99 (blue). (b) The amino acid residue root-mean-square-fluctuation (RMSF) of EspG_5mtu_ (starting structure PDB ID 4KXR, chain C) during a molecular dynamics simulation run. The secondary structure assignment of the crystallographic structure is shown above the RMSF plot for reference, where β-sheets and α-helices are represented by arrows and rectangles, respectively. (c) The flexible loop regions and the amino acid residues missing from the crystallographic structure of EspG_5mtu_ crystal structure when in complex with PE25-PPE41 (purple) are modeled as dummy residues (blue spheres).

Taken together, these data show clear differences in the oligomerization trends of each EspG ortholog from different ESX systems, indicating that this could be another level of system specificity involved in the secretion of PE–PPE proteins via ESX systems.

### Model of the EspG_3_-PE5–PPE4 trimer structure adopts an extended conformation

To further analyze substrate recognition by different EspG proteins, we measured solution scattering of EspG_3_–PE5–PPE4 and EspG_5_–PE25–PPE41 complexes from *M. tuberculosis*. Model-free parameters derived from the SAXS data indicate that EspG_3msm_–PE5–PPE4 has a more extended open overall conformation (*D_max_* =14.5 nm / *R_g_* = 4.25 nm) than the crystallographic structure of the EspG_5mtu_–PE25–PPE41 complex (*D_max_* = 13.0 nm / *R_g_* = 3.93 nm) (Fig. 6a). *Ab initio* modeling in *P*1 symmetry using SAXS data provided independent information about the overall shapes. The EspG_3msm_ complex reveals a more open structure than the EspG_5mtu_ complex, which is consistent with the model-free parameters. In addition, the overall shape of the EspG_5mtu_ complex *ab initio* structure is in agreement with the crystallographic structure showing a compact structure (PDB ID 4KXR; NSD = 1.7, data not shown). The theoretical scattering of the crystallographic structure of the EspG_5mtu_–PE25– PPE41 complex fits the measured SAXS data very well (χ^2^ = 0.95) (Fig. 6b). This indicates that, in solution, the EspG_5mtu_ β2–β3 loop is in the extended conformation seen in the crystal structure of the EspG_5mtu_–PE25–PPE41 heterotrimer. Thus, the SAXS data for EspG_5mtu_ alone and in complex with PE25–PPE41 indicates that the β2–β3 loop undergoes a significant conformational change upon binding.

**Figure 6.**
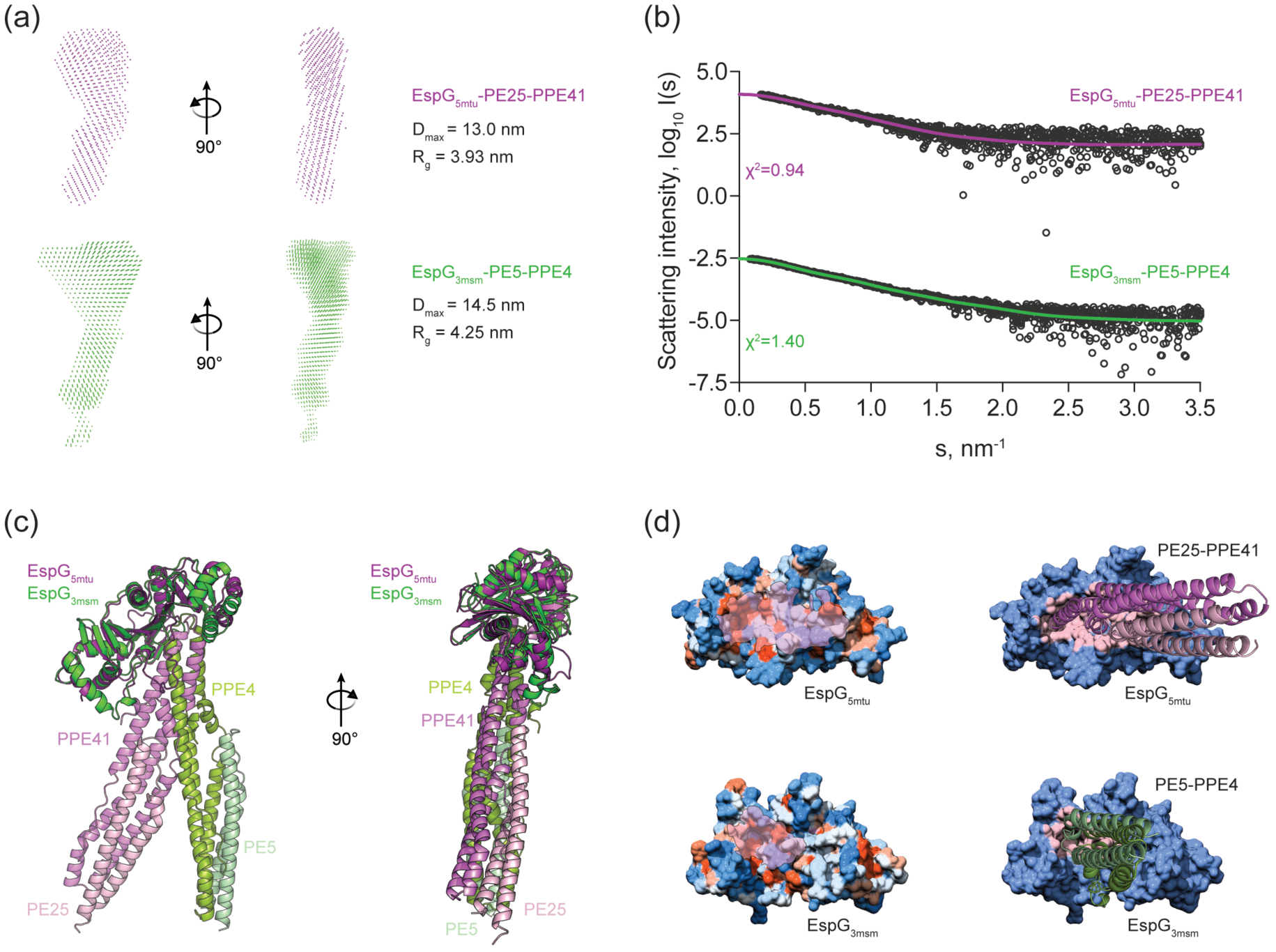
Comparison of EspG_5_-PE25-PPE41 and EspG_3_-PE5-PPE4^1-178^ complexes structures. (a) SAXS-based *ab initio* models of the EspG_5mtu_-PE25-PPE41 complex (purple) and the EspG_3msm_-PE5–PPE4^1-178^ complex (green) reveal a more extended conformation of the EspG_3msm_-PE5–PPE4^1-178^ complex, as indicated by the maximum diameter (*D_max_*) and radius of gyration (*R_g_*) of each complex. (b) Experimental SAXS data (black circles) with the computed fits (solid lines) and the respective discrepancy values (*χ^2^*) for EspG_5mtu_-PE25- PPE41 and EspG_3msm_-PE5-PPE4 protein complexes. The theoretical SAXS curve, calculated with CRYSOL from the EspG_5mtu_-PE25–PPE41 complex structure (PDB ID 4KXR), fits the experimental SAXS data with a goodness-of-fit (*χ^2^*) of 0.95 (purple). The theoretical SAXS curve calculated from the EspG_3msm_ rigid-body model in complex with a homology model of PE5-PPE4^1-178^ fits the experimental SAXS data (green) with a goodness-of-fit values (*χ^2^*) of 1.40. (c) Superposition of the crystal structure of EspG_5mtu_-PE25-PPE41 (purple-lilac-light pink) and the SAXS-derived rigid body model of EspG_3msm_-PE5-PPE4^1-178^ (dark green-light green-light blue). (d) Hydrophobicity surface representation of EspG proteins with PPE-interacting surface highlighted in transparent pink (left panel). Interaction interface of EspG_5mtu_-PPE41 and EspG_3msm_-PPE4, with PPE-PE proteins represented as cartoon and EspG proteins as surface with the contact residues in the interface colored in light pink in each EspG protein (right panel).

In order to produce an atomic rigid body model of the EspG_3msm_–PE5–PPE4 complex, 300 decoy structures were generated using a molecular docking approach [35] and the crystallographic EspG_3msm_ (PDB ID 4L4W, chain B) structure together with homology models of a complex of full length PE5 and the N-terminal domain of PPE4 (residues 1-178) from *M. smegmatis* generated with *SWISS-MODEL* [36]. The decoy structures were then ranked by the goodness-of-fit values of their theoretical scattering compared to the experimental scattering data. Fourteen complex structures were selected for further analysis on the basis of their chi^2^–values (χ^2^ < 1.7). In addition, all structures had acceptable fits according to *p*- values provided by the correlation map approach [37]. A representative model with the lowest χ^2^–value was selected for the comparison with the EspG_5mtu_–PE25–PPE41 complex (Fig. 6b-d).

Comparison of the crystallographic structure of EspG_5mtu_–PE25–PPE41 (PDB ID 4KXR, [26]) and the SAXS-based rigid-body model of EspG_3msm_–PE5–PPE4 shows similarities in the binding interfaces of the two complexes, but also significant differences related to the overall binding orientation (Fig. 6c). As expected, the interface between EspG_3msm_ and the PE5– PPE4 heterodimer was found to be mostly comprised of hydrophobic amino acid residues allowing interaction with the hh-motif of PPE4 [26] (Fig. 6d). Analogous to the EspG_5mtu_– PE25–PPE41 complex, the loop between helices α4 and α5 of PPE4 (Ala^125^-lIe^134^) interacts with the central β-sheet of EspG_3_. The structure suggests a hydrogen bonding network between EspG_3_ and PPE4 formed by several hydrophobic, polar and charged amino acid residues Trp27, Glu196, Glu211, Ser215 of EspG_3msm_ and residues Thr126, Phe128, Gly130, Asn132, Thr133, Ile134 of PPE4 from *M. smegmatis*. To validate our SAXS model and the interface of EspG_3msm_ and PPE4 in particular, we constructed two mutants of EspG_3msm_: E196R and S215Y. Substitution E196R reverses the charge of a conserved E196 residue in the EspG_3_ orthologs, whereas mutation S215Y introduces a bulky residue that would sterically prevent binding of PPE4. Pull-down experiments showed that E196R and S215Y variants of EspG_3msm_ could not bind PE5–PPE4 dimers (Supplementary Fig. S8).

Based on the SAXS data and models, we suggest that the more open and flexible structure of EspG_3_–PE5–PPE4 is due to differences in the EspG_3_ β2–β3 loop region. The EspG_5mtu_ β2–β3 loop is comprised of 23 amino acid residues (Gly^92^-Asn^114^) and its interaction with PE25–PPE41 forms an extended interface extending from the central β-sheet of EspG_5mtu_. However, the homologous loop region in the EspG_3msm_ structure is significantly shorter (12 amino acid residues; Val^87^ to Leu^98^) thus an analogous interaction is missing in the EspG_3msm_–PE5–PPE4 rigid-body model. Given that ESX-5 is the major secretion system for export of PE/PPE proteins [22–24, 32], the length and flexibility of the EspG_5mtu_ β2-β3 loop could be an important structural feature allowing EspG_5mtu_ to interact specifically with many different PE–PPE heterodimers.

## Conclusions

Transport of multiple proteins across the mycobacterial cell envelope is facilitated by the ESX system and cytoplasmic EspG chaperones [38]. However, the precise recognition mechanism of the cognate substrates by ESX system-specific chaperones is not yet fully understood. In this work we present structural analyses of EspG_1_, EspG_3_ and EspG_5_ and their complexes with PE–PPE secretion substrates using a combination of experimental and *in silico* methods. The solution scattering data together with novel X-ray crystallographic structures allows us to hypothesize about the substrate specificity of EspG chaperones and provides insight into chaperone-substrate binding mechanisms. This study demonstrates that the β2–β3 loop of EspG plays an important role in PE–PPE binding and is a major differentiation factor between the EspG chaperones of orthologous ESX secretion systems. Our results also suggest that EspG dimerization may play a role in substrate recognition.

## Materials and methods

### Cloning, expression and purification of *M. kansasii* EspG_1_

The DNA sequence corresponding to the full-length EspG_1mka_ was PCR amplified using primers G1mka_F1Nde, 5’-GATACATATGACCGGTCCGCTCGCTAC and G1mka_R283Hind, 5’-CTCAAGCTTAGCCTCGGGCGGAGGCTTG, and genomic DNA of *M. kansasii* ATCC 12478. The PCR product was digested with NdeI/HindIII and ligated into the corresponding sites of a modified pET-28b vector to create an N-terminal His_6_-tag with a tobacco etch virus (TEV) cleavage site. In efforts to optimize initial crystals, the Cys114Ala and Cys170Ala mutations were introduced using the QuikChange protocol (Stratagene). A truncated DNA fragment corresponding to residues 17-271 was PCR amplified using primers G1mka_F17Nco, 5’-GATTCCATGGTCGGCGTCGAGGTCACC and G1mka_R271Hind, 5’-CTCAAGCTTCAATCTAACCAGGAGCCCGC and cloned into a pET-based vector containing an N-terminal His_6_-tag followed by a TEV cleavage site and T4L sequence (residues 2-162). The T4L sequence corresponds to a cysteine-less variant with Cys54Thr and Cys97Ala mutations [39, 40]. T4L-EspG_1mka_ was expressed in *E*. *coli* Rosetta2(DE3) cells in LB supplemented with 50 μg mL^-1^ kanamycin and 34 μg mL^-1^ chloramphenicol. Cells were grown at 37°C and expression was induced with 0.5 mM IPTG at *A*_600_ of 0.6. Cells were harvested by centrifugation after 3 h, resuspended in lysis buffer (20 mM Tris-HCl pH 8.5, 300 mM NaCl, 10 mM imidazole) and lysed using an EmulsiFlex C5 homogenizer (Avestin). EspG_3msm_ was purified via Ni-NTA metal affinity chromatography. The His_6_-tag was cleaved using TEV protease followed by a second Ni-NTA purification step to remove uncleaved T4L-EspG_1mka_ and His_6_-tagged TEV protease. Size-exclusion chromatography was performed using a Superdex 200 column (GE Biosciences) equilibrated in buffer containing 20 mM HEPES pH 7.5, 300 mM NaCl.

### Crystallization and structure solution of T4L-EspG_1mka_

Crystals of T4L-EspG_1mka_ were obtained by the hanging drop vapor diffusion method using crystallization solution containing 0.1 M Bicine pH 9.0, 1.0 M LiCl, 10% PEG6000. The crystals were cryoprotected in crystallization solution supplemented with 20% glycerol and flash cooled in liquid nitrogen before data collection. Data were collected at the SER-CAT beamline 22-ID at the Advanced Photon Source, Argonne National Laboratory. Data were processed and scaled using *XDS* and *XSCALE* [41]. CC_1/2_ value of 0.5 for the outer shell was used to determine the resolution [42]. The T4L-EspG_1mka_ structure was solved by molecular replacement using *Phaser* [43], with the T4L structure (PDB 4GBR) [44] and poly-Ala EspG_3msm_ (PDB ID 4L4W) structure as search models. Two copies of T4L-EspG_1mka_ were located in the asymmetric unit. Density modification was performed using *Parrot* [45], and the molecular replacement model was re-built using *Buccaneer* [46, 47], *ARP/wARP* [48] and manual building in *Coot* [49]. The iterative rounds of refinement and re-building were performed using *phenix.refine* [50] and *Coot*. Non-crystallographic symmetry (NCS) restraints were applied throughout the refinement. Statistics for data collection, refinement, and model quality are listed in Table 1.

### Cloning, expression, and purification of *M. marinum* EspG_3_

The gene (MMAR_0548) encoding EspG_3mma_ was PCR-amplified from *M*. *marinum* M genomic DNA with Phusion DNA polymerase (New England Biolabs) using gene specific primers (MMAR_0548.For, 5’-AACCTGTATTTCCAGAGTATGGAGTCAATGCCCAACG and MMAR_0548.Rev, 5’-ttcgggctttgttagcagttaGGAGGGTTGACTCGAGAAATCT) and was cloned into a modified pET28 vector, pMAPLe4 [51], using the Gibson ISO assembly procedure [52]. The DNA sequence of the construct was verified by DNA sequencing (Genewiz). EspG_3mma_ was expressed from pMAPLe4 as a maltose binding protein (MBP) fusion which was cleaved *in vivo* from MBP via a tobacco vein mottling virus (TVMV) protease cleavage site situated between the two moieties; a His_6_ affinity tag and a tobacco etch virus (TEV) protease cleavage site is encoded in the linker between the TVMV protease cleavage site and the N-terminus of the target protein. EspG_3mma_ was expressed in *E*. *coli* BL21-Gold (DE3) (Agilent Technologies) using Terrific broth media and protein expression was induced with 1 mM IPTG at 18°C overnight. Harvested cells were resuspended in lysis buffer (20 mM Tris, pH 8.0, 300 mM NaCl, 10% glycerol, 10 mM imidazole) supplemented with β-mercaptoethanol (2 mM), DNase I, lysozyme, and Complete protease inhibitor cocktail (Roche) and lysed by sonication. The lysate was centrifuged (39,000 *g*, 30 minutes, 4°C) and EspG_3mma_ was purified from the clarified supernatant using Ni-NTA resin (Thermo Fisher Scientific) equilibrated in lysis buffer. The bound protein was eluted with lysis buffer containing 300 mM imidazole and further purified by size exclusion chromatography (SEC) using a HiLoad 16/60 Superdex 75 column (GE Healthcare) equilibrated with 20 mM Tris, pH 8.0, 300 mM NaCl, 10% glycerol. EspG_3mma_ eluted from the column in a single symmetrical peak which was concentrated to 21.5 mg mL^-1^ for crystallization screening.

### Crystallization and structure determination of EspG_3mma_

Small crystals of EspG_3mma_ were grown using the hanging drop vapor-diffusion method by mixing 1 μL of protein with 0.5 μL of reservoir solution (1.4 M ammonium sulfate, 200 mM lithium sulfate, 60 mM CAPS, pH 10.5). These small crystals were used to streak seed other drops that had been equilibrated for a week but showed no signs of crystal growth. Large crystals were found in seeded drops with a reservoir solution containing 1.45 M ammonium sulfate, 200 mM lithium sulfate, 70 mM CAPS, pH 10.5 after 5 weeks. For phase determination and cryoprotection crystals were soaked for 30 minutes at room temperature in a solution containing approximately 4 mM platinum potassium thiocyanate, 1.17 M ammonium sulfate, 160 mM lithium sulfate, 56 mM CAPS pH 10.5, and 15.5% glycerol. Diffraction data were collected at the Advanced Photon Source at Argonne National Laboratory on beamline 24-ID-C. The data were processed with *XDS* [41], and the structure was solved by single wavelength anomalous dispersion using *HKL2MAP* [53], in the *SHELX* suite of programs [54], which determined the position of seven platinum atoms in the K_2_Pt(SCN)_6_–soaked crystal. An initial model was built using *SHELXE* [55] which was improved through iterative rounds of manual model building using *Coot* [49] interspersed with refinement using *REFMAC*5 [56].

### Cloning, expression and purification of *M. smegmatis* EspG_3_

The gene *msmeg_0622* encoding EspG_3msm_ was PCR amplified from genomic DNA of *M. smegmatis* mc^2^155 using primers MsmG3_F1Nde, 5’- GAGACATATGGGGCCTAACGCTGTTG, and MsmG3_R293Hind, 5’-CTCAAGCTTACTAGTCATGCTTTCTGGGTTCTTCTCTG. The PCR product was digested with NdeI/HindIII and ligated into the corresponding sites of a modified pET-28b vector to create TEV protease-cleavable N-terminal His_6_-tag fusion. The construct was verified by DNA sequencing (Eurofins Genomics). EspG_3msm_ was expressed and purified using procedures similar to the T4L-EspG_1mka_ fusion, except the final SEC step was performed in buffer containing 20 mM HEPES pH 7.5, 100 mM NaCl.

### Crystallization and structure solution of EspG_3msm_

Crystals of EspG_3msm_ in space group *P*3_2_21 were obtained by hanging drop vapor diffusion method with crystallization solutions containing 0.1 M Tris-HCl pH 7.0, 0.2 M Mg acetate, 1.8 M NaCl (SeMet substituted EspG_3msm_) and 0.1 M HEPES pH 7.4, 1.0 M LiCl, 10% PEG6000 (native EspG_3msm_). Crystals were transferred to crystallization solutions supplemented with 20-25% glycerol and flash-cooled in liquid nitrogen prior to data collection. Data for native and SeMet substituted EspG_3msm_ crystals were collected at the SER-CAT beamline 22-ID at the Advanced Photon Source, Argonne National Laboratory. Data were processed and scaled using *XDS* and *XSCALE* [41]. The EspG_3msm_ structure was solved by SeMet-SAD. The initial selenium positions were found with *SHELXD* [54] using *HKL2MAP* interface [53]. Phasing, density modification and initial model building was performed using autoSHARP [57]. A partial model was refined against a native dataset and rebuilt using *REFMAC*5 [56], *ARP/wARP* [48] and AutoBuild within *PHENIX* [58]. The model was completed by iterative rounds of refinement and rebuilding using *phenix.refine* [50] and *Coot* [49].

Crystals of SeMet substituted EspG_3msm_ in space group *C*222_1_ were grown using crystallization solution containing 0.1 M Tris-HCl pH 8.5, 1.0 M LiCl, 20% PEG6000. Crystals were transferred into crystallization solution supplemented with 20% glycerol and flash cooled in liquid nitrogen. The structure was solved by molecular replacement using *Phaser* [43] with EspG_3msm_ (PDB ID 5SXL) structure as a search model. Two EspG_3msm_ molecules were located in the asymmetric unit. Following density modification using *Parrot* [45], the molecular replacement model was rebuilt using *Buccaneer* [46, 47]. The model was further improved using *Coot*, *ARP/wARP* and AutoBuild within *PHENIX*. The final model was refined using *phenix.refine*.

Crystals of EspG_3msm_ in space group *P*4_3_2_1_2 were grown using crystallization solution containing 0.1 M Na cacodylate pH 6.0, 15% PEG200, 5% PEG3350. The crystals were cryoprotected in solution containing 0.1 M Na cacodylate pH 6.0, 35% PEG200, 5% PEG3350 and flash cooled in liquid nitrogen. Molecular replacement using *Phaser* and EspG_3msm_ (PDB ID 4L4W) structure as a search model located 2 molecules in the asymmetric unit. The model was refined using *REFMAC*5 and rebuilt using AutoBuild within *PHENIX*. The iterative rounds of refinement and rebuilding were performed using *phenix.refine* and *Coot*. NCS restraints were applied in early rounds of refinement and were later omitted as the model quality improved. The last several rounds of refinement were performed using 4 translation/libration/screw (TLS) groups, identified by the TLSMD server [59, 60], per protein chain.

### Sample preparation for SAXS measurements

The gene MMAR_5441 encoding EspG_1mma_ was PCR amplified from genomic DNA of *M. marinum E11* using primers DF018, 5’- ATATATAGATCTACCGGTCCGCTCGCTACCGG’, and DF019, 5’- ATATATATGCGGCCGCTTAACCTCGGGCGGTGGCGTCG’. The PCR product was digested with BglII/NotI. The gene MMAR_0548 encoding EspG_3mma_ was PCR amplified from genomic DNA of *M. marinum E11* using primers DF020, 5’- ATATATACCGGTGGAATGGAGTCAATGCCCAACGC’, and DF021, 5’- ATATATATGCGGCCGCTTAGGAGGGTTGACTCGAGAA’. The PCR product was digested with AgeI/NotI. Clones containing genes Rv0289 and Rv1794 encoding EspG_3mtu_ and EspG_5mtu_ were further digested with BglII/NotI and AgeI/NotI. Digested fragments were ligated into the corresponding sites of pETM11-SUMO3 to create SENP-2 protease-cleavable N-terminal His_6_-SUMO_3_-tag fusions.

His_6_-SUMO_3_-EspG_1mma_, His_6_-SUMO_3_-EspG_3mma_, His_6_-SUMO_3_-EspG_3mtu_ and His_6_-SUMO_3_-EspG_5mtu_ proteins for SAXS measurements were purified as follows: cells were resuspended in lysis buffer 20 mM HEPES pH 7.5, 150 mM NaCl, 10 mM imidazole, 0.25 mM tris(2-carboxyethyl)phosphine (TCEP), 10% (w/v) glycerol (pH 7.5) containing 1/100 protease inhibitor mix HP (Serva), DNAse I (10 μg/ml) and disrupted by lysozyme treatment followed by sonication. The protein was purified via Ni-NTA (Qiagen) affinity chromatography. His6-SUMO_3_-tag was cleaved by SenP_2_ protease and further purified using a Phenyl Sepharose HP column (GE Biosciences), followed by SEC using a Superdex 200 16/60 column (GE Biosciences) pre-equilibrated with 20 mM HEPES pH 7.5, 150 mM NaCl, 0.25 mM TCEP, 10% (w/v) glycerol.

The purification procedure for the EspG_5mtu_-PE25-PPE41-His_6_ complex was the same as for the His_6_-SUMO_3_-EspG proteins described above with the exception that the cleavage of the His6 tag was performed by addition of TEV protease. For final aggregated protein removal, the complex was concentrated and injected into a Superdex 200 16/60 size-exclusion chromatography column (GE Biosciences) pre-equilibrated with 20 mM HEPES pH 7.5, 150 mM NaCl, 0.25 mM TCEP, 10% (w/v) glycerol. EspG_3msm_–PE5–PPE4 complex was obtained as described in [26], and further purified using a Superdex 200 column equilibrated in buffer containing 20 mM HEPES pH 7.5, 100 mM NaCl. All samples used for SAXS experiments were concentrated to the appropriate protein concentrations ranging from ~1–7 mg ml^-1^.

Mutations E196R and S215Y were introduced into EspG_3msm_ using Gibson mutagenesis protocol and following primers: G3msm_E196R_F 5’-GGTCGGCGCACCTACGTCCGTATCGTCGCGGGCGAGCAT, G3msm_E196R_R 5’-ATGCTCGCCCGCGACGATACGGACGTAGGTGCGCCGACC, G3msm_S215Y_F 5’-CACCACCGAGGTGGGGGTCTACATCATCGACACCCCACAC, G3msm_S215Y_R 5’-GTGTGGGGTGTCGATGATGTAGACCCCCACCTCGGTGGTG, IsoKan_1 5’-GACAATTACAAACAGGAATCGAATGC and IsoKan_2 5’-GCATTCGATTCCTGTTTGTAATTGTC. The pull-down experiments were performed as described in [26].

### SAXS measurements

SAXS measurements were carried out at beamline P12 (EMBL/DESY, Hamburg) [61] at the PETRA-III storage ring using a Pilatus 2M detector (Dectris). Measurements for the purified proteins were made at several concentrations (Table 4). For each measurement twenty 50 ms exposure frames were collected and averaged using a sample volume of 30 μl at a temperature of 10°C. The SAXS camera was set to a sample-detector distance of 3.1 m, covering the momentum transfer range 0.008 Å-1 < *s* < 0.47 Å^-1^ (*s* = 4π sin(θ)/λ where *2θ* is the scattering angle and λ=1.24 Å is the X-ray wavelength). Prior to and following each sample exposure, a buffer control was measured to allow for background subtraction.

### SAXS data analysis using model-free parameters

Radius of gyration *R_g_* and forward scattering intensity *I*(0), were independently determined using Guinier analysis [62] and the indirect Fourier transformation approach of the program GNOM [63]. Additionally, the maximum particle dimension *D_max_* was obtained from the latter approach. Molecular masses of protein constructs (MM_SAXS_) were calculated by comparing the extrapolated forward scattering intensities with that of a reference BSA sample (MM_ref_ = 66 kDa) together with concentration information. The excluded volume of the hydrated protein *Vp* was obtained with DATPOROD [64] and used to extract an independent estimate of molecular mass (MM_POROD_). For globular proteins, hydrated protein volumes in Å^3^ are approximately 1.7 times the molecular masses in Dalton.

### SAXS-based structural modeling

*Ab initio* models were reconstructed from the scattering data using the simulated annealing based bead-modeling program *DAMMIF* [65]. Ten independent reconstructions were averaged to generate a representative model with the program *DAMAVER* [66]. In addition, the average *DAMMIF ab initio* model was used to calculate an excluded volume of the particle, *V_DAM_*, from which an additional independent MM estimate can be derived (empirically, *MM_DAM_* ~ *V_DAM_/2*). The resolutions of the model ensembles were estimated by a Fourier Shell correlation approach [67].

Theoretical scattering profiles from available high-resolution crystallographic structural models were calculated using the program *CRYSOL* [68] and used to determine the fit of these models to the experimental scattering data. Given the atomic coordinates of a structural model, *CRYSOL* minimizes the discrepancy between the experimental and theoretical scattering intensities by adjusting the excluded volume of the particle and the contrast of the hydration layer. Rigid-body modeling was performed using the *ZDOCK* docking approach [35] to generate decoy structures and complexes were ranked based on their fit to the experimental scattering data.

The scattering profile from a molecular mixture can be decomposed into a linear combination of individual contributions *I_i_*(*s*) from the different species. If the structures of the components are known or their individual scattering profiles can be measured, the volume fractions of the species that fit the SAXS data can be determined by the program *OLIGOMER* utilizing nonnegative least-squares fitting [69]. Dimeric structures of EspG_3_ from *M*. *tuberculosis* and *M*. *marinum* were generated from their monomeric crystallographic structures (PDB IDs 4W4I and 5DBL, respectively) using the *M. smegmatis* EspG_3_ dimer structure (PDB ID 4L4W) as an interaction template. We also used the program *OLIGOMER* [69] for fitting of experimental scattering profiles of EspG_3mtu_ and EspG_3mma_ by weighted combinations of theoretical scattering profiles from the monomeric crystallographic structures and dimeric models. In the case of EspG_3msm_, theoretical scattering profiles based on the dimeric and monomeric crystallographic structures (PDB ID 4L4W) were used as inputs for *OLIGOMER*. For EspG_1mma_, two different dimeric structures were tested: the first dimer structure is based on the structure of the EspG_1mka_ dimer (PDB ID 5VBA) while the second model for a EspG_1mma_ dimer was constructed using the EspG_3msm_ dimer structure (PDB ID 4L4W) as a template.

Flexibility analyses of protein structures in solution were conducted using their crystallographic structures as starting points for the ensemble optimization method (EOM). This approach seeks to best fit the experimental scattering profile with an ensemble of conformations [70, 71]. Possible conformations of loop regions were modeled with the program *RANCH* producing 10,000 random configurations, while the rest of the protein was kept fixed. A genetic algorithm was employed to find the set of conformations best fitting the SAXS data. The structures selected from the random pool of structures were analyzed with respect to the *R_g_* distribution.

### Molecular dynamics simulations

The program NAMD was employed with the CHARMM27 force field for description of the protein and the TIP3P solvent model for water. Constant particle number, constant pressure and constant temperature (*NpT*) ensembles were assumed [72–74]. *Langevin* dynamics were used to maintain constant temperature. Pressure was controlled using a hybrid *Nose*-*Hoover Langevin* piston method. An in-house computational pipeline for high-throughput MD simulations and the visualization program *VMD* were used to prepare input files and to analyze the simulation trajectories [75].

### Accession numbers

The structure factors and atomic coordinates have been deposited in the Protein Data Bank under accession codes 5VBA (T4L-EspG_1mka_), 5DLB (EspG_3mma_), 4L4W, 4RCL and 5SXL (EspG_3msm_). The SAXS data were deposited in the SASBDB under accession codes SASDDQ2, SASDDR2, SASDDS2, SASDDT2, SASDDU2, SASDDV2, SASDDW2, SASDDX2.

## Acknowledgements

We thank Marcel Behr for providing *M. kansasii* genomic DNA, Wilbert Bitter for providing *M. marinum* genomic DNA and Carlo Carolis for the *espG_3mtu_* construct. We thank the staff of the UCLA-DOE Institute Protein Expression Technology Center, supported by the U.S. Department of Energy, Office of Biological and Environmental Research (BER) program under Award Number DE-FC02-02ER63421, and the UCLA Crystallization Core for assistance in protein purification and crystallization screening. Authors thank staff members of the Northeastern Collaborative Access Team (NE-CAT) and Southeast Regional Collaborative Access Team (SER-CAT) at the Advanced Photon Source, Argonne National Laboratory, for assistance during data collection. Use of the Advanced Photon Source was supported by the U. S. Department of Energy, Office of Science, Office of Basic Energy Sciences, under Contract No. W-31-109-Eng-38. We acknowledge the Sample Preparation and Characterization (SPC) facility of EMBL at PETRA3 (DESY, Hamburg) for technical support. Work performed in the laboratory of D.E. is supported by the Howard Hughes Medical Institute and National Institutes of Health grants 23616-002-06 F3:02, TBSGC P01 (AI068135), and TBSGC P01 (AI095208). Research reported in this publication was partially supported by an Institutional Development Award (IDeA) from the National Institute of General Medical Sciences of the National Institutes of Health under grant numbers P20GM103486 and P30GM110787, and by the National Institute of Allergy and Infectious Diseases grant number R01AI119022 to KVK. A.T.T. was supported by the EMBL EIPOD program under Marie Curie COFUND actions and by the Bundesministerium für Bildung and Forschung (BMBF) project BIOSCAT (grant 05K12YE1).

## Abbreviations used

EOM: ensemble optimization method
MD: molecular dynamics
RMSD: root-mean-square deviation
NSD: normalized spatial discrepancy
RMSF: root-mean-square-fluctuations
SAXS: small-angle X-ray scattering
SEC: size-exclusion chromatography
SeMet: selenomethionine
TEV: tobacco etch virus
TCEP: tris(2-carboxyethyl)phosphine

